## Supplementary Figures for "Evaluating transportability of *in-vitro* cellular models to *in-vivo* human phenotypes using gene perturbation data"

Supplementary Figures 1-9

**Supplementary Figure 1** Intracellular insulin content and Hba1c

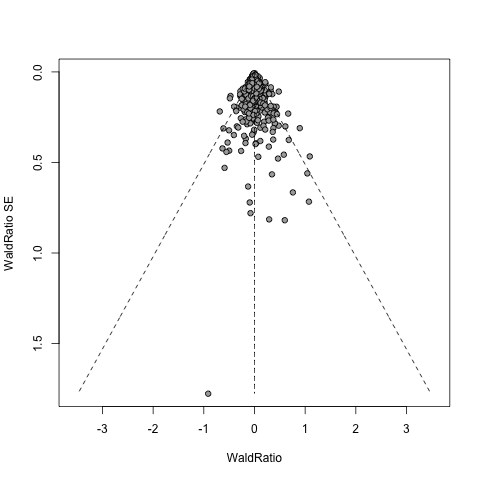

**Supplementary Figure 2** Intracellular insulin content and Type II diabetes

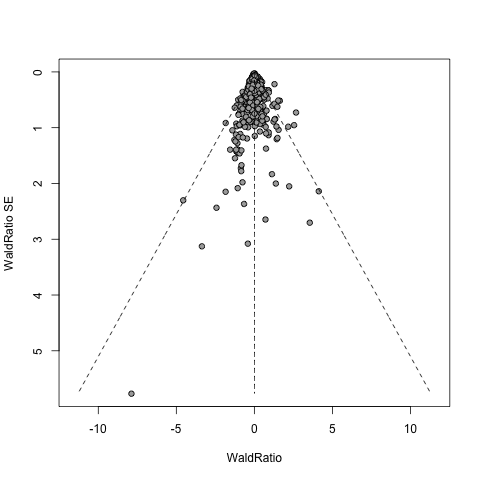

**Supplementary Figure 3** Adipocyte differentiation and body-mass index

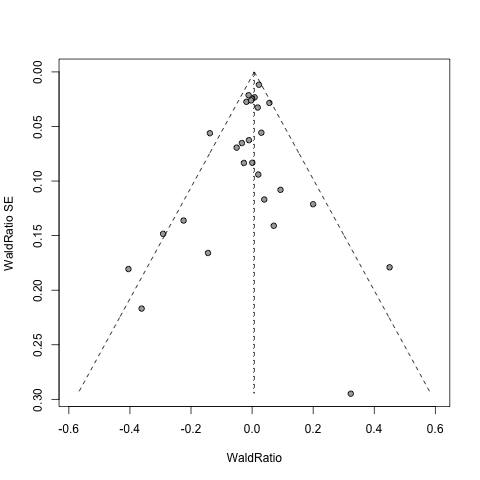

**Supplementary Figure 4** Adipocyte differentiation and waist-circumference

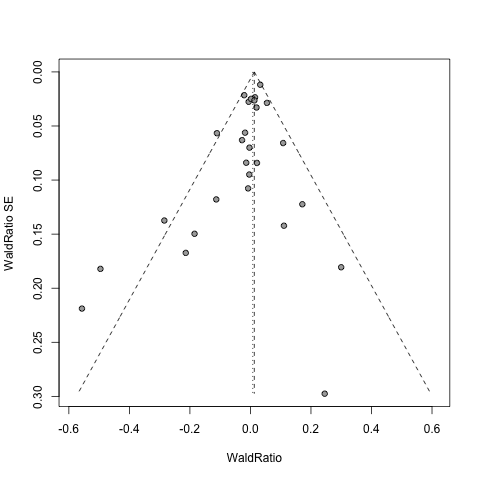

**Supplementary Figure 5** Adipocyte differentiation and body fat percentage

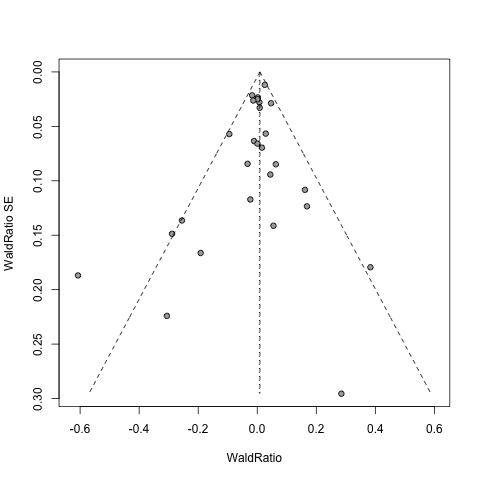

**Supplementary Figure 6** LoF-IV simulations: example of baseline model

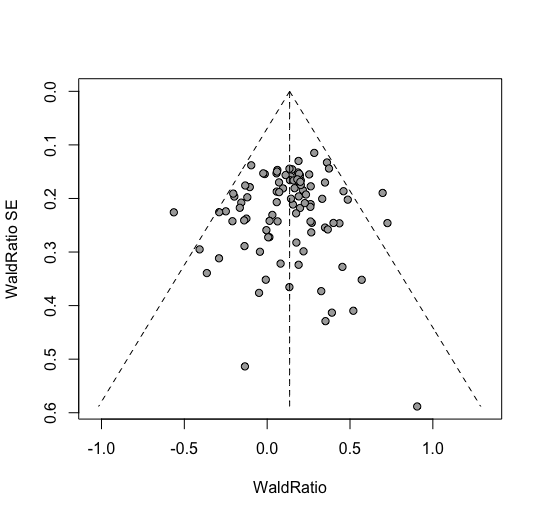

**Supplementary Figure 7** LoF-IV simulations: example of balanced pleiotropy model

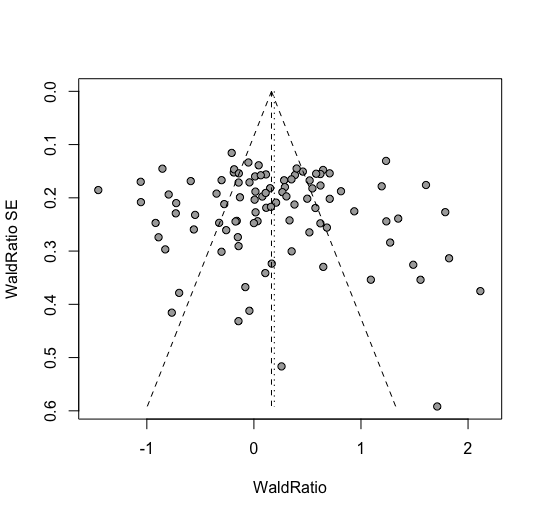

**Supplementary Figure 8** LoF-IV simulations: example of unbalanced pleiotropy model

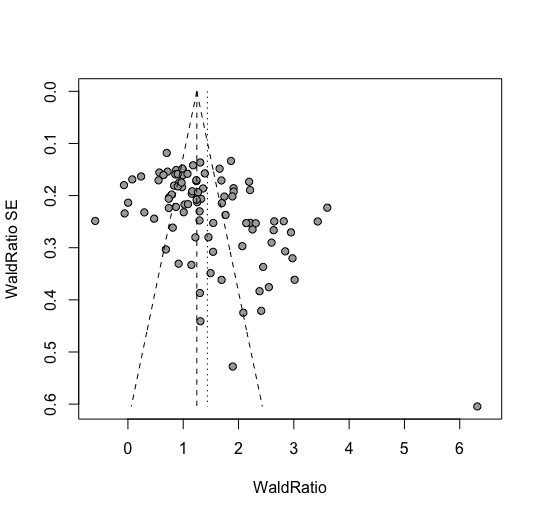

**Supplementary Figure 9** LoF-IV simulations: example of phenotypic pleiotropy model

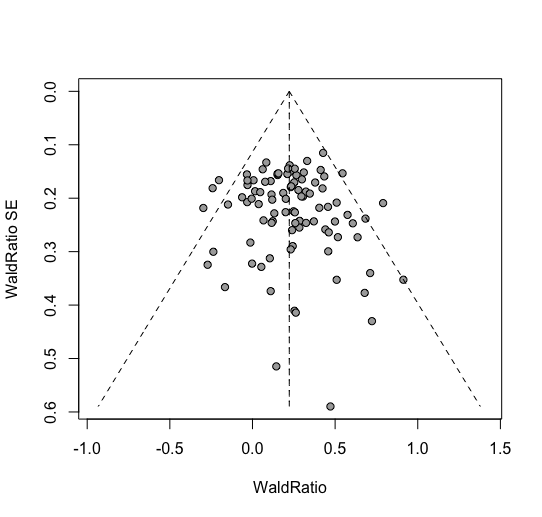
